# Viral enumeration using cost-effective wet-mount epifluorescence microscopy for aquatic ecosystems and modern microbialites

**DOI:** 10.1101/2023.07.07.548154

**Authors:** Madeline Bellanger, Pieter Visscher, Richard Allen White

## Abstract

Enumeration is a fundamental measure of community ecology in which viruses represent the most numerous biological identities. Epifluorescence microscopy (EFM) has been the gold standard method for environmental viral enumeration for over 25 years. Currently, standard EFM methods using the Anodisc filters are no longer cost-effective (>$15 per slide) and have yet to be applied to modern microbialites. Microbialites are microbially driven benthic organosedimentary deposits that have been present for most of Earth’s history. We present a cost-effective method for environmental viral enumeration from aquatic samples, microbial mats, and exopolymeric substances (EPS) within modern microbialites using EFM. Our integrated approach, which includes filtration, differential centrifugation, chloroform treatment, glutaraldehyde fixation, benzonase nuclease treatment, probe sonication (EPS and mat only), SYBR Gold staining, wet-mounting, and imaging, provides a robust method for modern microbialites and aquatic samples. Viral abundances of modern microbialites and aquatic samples collected from Fayetteville Green Lake (FGL) and Great Salt Lake (GSL) did not differ across ecosystems by sample type. EPS and microbial mat samples had an order of magnitude higher viral-like particle (VLP) abundance when compared to water regardless of the ecosystem (10^7^ vs. 10^6^). Viral enumeration allows for estimates of total viral numbers and weights. The entire weight of all the viruses in FGL and GSL are ∼598 g and ∼2.2 kg, respectively; this is equivalent to a loaf of bread for FGL and standard brick for GSL. Further development of EFM methods and software is needed for viral enumeration. Our method provides a robust and cost-effective (∼$0.75 per sample) viral enumeration within modern microbialites and aquatic ecosystems.

## 1. Introduction

Viruses are the most numerous ’biological entity’ on Earth, with a ubiquitous range across every environment and an estimated global abundance of 10^31^ viral-like particles (VLPs) (Fuhrman et al., 1999; Hendrix et al., 1999; Mushegian et al., 2020). This 10^31^ VLPs estimate (i.e., Hendrix product) accounts mainly for double-stranded DNA phage abundances (Hendrix et al., 1999; Mushegian et al., 2020). This estimation did not include the diversity of large DNA, RNA, and ssDNA (White III et al., 2021). Viral abundance estimates may be an underestimation as 10^30^ viruses are estimated to exist in the ocean alone (Suttle et al., 2007), and global biomass estimates (Bar-On et al., 2018) suggest more measurements and method developments are still needed.

Epifluorescence microscopy (EFM) is the gold standard technique for the enumeration of microbes and viruses (Noble & Fuhrman, 1998), which includes the enumeration of aquatic bacteria since 1977 (Hobbie et al., 1977), but its use for viruses began in the early 1990s (Hara et al., 1991). Viruses play critical roles in the life and death cycles of microbes through viral lysis that interplays in global biogeochemical cycles (e.g., carbon and nitrogen pool), population dynamics, the marine viral shunt, and nutrient transportation (Wilhelm & Suttle, 1999; Weinbauer, 2004; Suttle, 2007; Rohwer et al., 2009; Jiao & Zheng, 2011; Breitbart, 2012; Lara et al., 2017; Wang et al., 2022). Aquatic systems remain the most enumerated ecosystems for viruses, with soils and sediments being a close second, and microbial mats rarely, followed by none for modern microbialites. Enumeration is the first critical step in understanding an ecosystem; thus, viral enumeration represents a major factor in microbial community ecology.

Microbial mats are laminated organosedimentary ecosystems that can form microbialite carbonate structures through mineral precipitation (i.e., lithification, Dupraz & Visscher, 2005; Dupraz et al., 2009; 2011). Microbialites are benthic microbial deposits (Burne & Moore, 1987) that accrete as a result of a microbial community trapping and binding sediments and by forming the locus of mineral precipitation (Dupraz & Visscher, 2005; Dupraz et al., 2009; 2011). Not all microbial mats can become microbialites; this depends both on abiotic (e.g., water hardness, cation concentration) and biotic (e.g., microbial metabolic activity) (White III et al., 2020). Microbialites have been present for 80% of geological history (e.g., 3.7 Gyr) and provide a modern proxy for ancient ecosystem formation (Nutman et al., 2016; White III et al., 2020). Precipitation of carbonates is facilitated by cyanobacterial photosynthetic alkalization and filament trapping/binding using exopolymeric substances (EPS) (Dupraz & Visscher, 2005; Dupraz et al., 2009; 2011). EPS is primarily produced by cyanobacteria and some heterotrophic bacteria within mats and trap sediments, allowing for the stabilization and accretion of carbonates (Dupraz & Visscher, 2005; Dupraz et al., 2009; 2011). EPS density, pH, and carbohydrate structure act as the site of mineral nucleation by binding cations that can influence mineral morphologies and carbonate precipitation (Decho, 1990; Braissant et al., 2003). EPS has also been shown to bind viruses and microbes within microbial mats (Carreira et al., 2015). Viruses, particularly cyanophages, have been proposed as the missing mechanism for understanding how soft microbial mats transition to hard carbonate microbialites (White III et al., 2021). Virus-host interactions may support the formation of microbialites through the alteration of microbial metabolism (specifically photosynthesis), the development of viral resistance strategies, or gene alteration impacting EPS production (White III et al., 2021).

EFM and flow cytometry (FCM) are the most widely utilized methods for enumerating VLPs (Noble & Fuhrman, 1998; Marie et al., 1999; Brussaard, 2004). Classically, viral enumeration was measured by transmission electron microscopy (TEM) (Weinbauer & Suttle, 1997; Noble & Fuhrman, 1998; Patel et al., 2007). TEM approaches for viral enumeration were problematic for routine use due to it being labor-intensive, having large specimen variability, and being expensive (Weinbauer & Suttle, 1997; Noble & Fuhrman, 1998; Patel et al., 2007). EFM was more accurate and precise in measuring viruses than TEM (Weinbauer & Suttle, 1997). Current EFM viral enumeration methods require expensive Whatman Anodisc filter 0.02 μm (catalog number WHA68096002) membranes (i.e., $16.32 per filter). These filters are required for EFM enumeration for viruses. However, they are only produced by a single manufacturer and have had shifts in supply and price increases via global inflation due to the COVID-19 pandemic. The filters have been over $10 USD for over a decade with no sign of price reduction. A more cost-effective method is needed to scale global sampling enumeration for viruses.

EFM viral enumeration has been previously applied to microbial mats but not microbialites (Carreira et al., 2015). Carreira et al. (2015) tested a variety of buffers (e.g., tetrasodium pyrophosphate vs. EDTA), nucleases (e.g., DNase I/RNase I vs. benzonase), and sonication methods (e.g., bath vs. probe). The Carreira et al. (2015) method suggested optimal EFM conditions for microbial mats: glutaraldehyde fixation (i.e., 2%), 0.1 mM EDTA, and benzonase treatment with probe sonication. However, this method requires Anodisc membrane filters, which makes it no longer cost-effective.

Wet-mount EFM has been suggested as a cost-effective alternative to Whatman Anodisc filter membrane-based protocols (Cunningham et al., 2015). This EFM method relies on chemical flocculation concentration of the VLPs using iron chloride precipitation, followed by EDTA-ascorbate resuspension, then SYBR gold nucleic acid staining, and wet-mounting on a standard glass slide (Cunningham et al., 2015). With and without chemical flocculation-based wet mount, EFM showed similar concordance and precision with Anodisc-based methods at a much lower cost (Cunningham et al., 2015). It also was effective for natural ambient marine and freshwater ecosystems from the low end of 1 × 10^6^ ml^-1^ to a high end of 1 × 10^8^ ml^-1^, similar to Anodisc methods (Cunningham et al., 2015). The wet-mount method has yet to be applied for solid non-aquatic samples such as soils, microbial mats, sediments, or microbialites.

There has been an ongoing debate about whether membrane-derived vesicles (MDVs), free extracellular DNA (FED), gene transfer agents, and cell debris may produce ‘fake particles’ that are labeled as VLPs (Forterre et al., 2013). They suggested improving EFM and FCM for viral enumeration by adding a chloroform step to limit MDVs and cellular debris and a nuclease step to remove FED (Forterre et al., 2013). Ribosomes with rRNAs could also be co-purified in viral preparations, which could be mistaken for viruses if clumped together but are reduced by chloroform treatment (Conceição-Neto et al., 2015). Membrane-bound viruses are sensitive to chloroform and some phage (e.g., *Corticoviridae* and *Inoviridae*) (Forterre et al., 2013). Viruses can be resistant to nuclease treatment, including RNA phages (e.g., MS2) (Carreira et al., 2015; Mikel et al., 2015)

Here we present a cost-effective method of EFM to enumerate viruses in aquatic environments, microbial mats, EPS within microbial mats, and modern microbialites within the Great Salt Lake (GSL) and Fayetteville Green Lake (FGL). Our method incorporates wet-mount (Cunningham et al., 2015), Carreira et al. (2015), and the suggestions of Forterre et al. (2013) to have a uniform method of EFM across aquatic ecosystems, microbial mats, and microbialites. Enumeration of viruses within modern microbialites could illuminate the microbial-viral-mineral interactions and mechanisms required to transition from soft microbial mat to hard carbonate lithified microbialite.

## 2. Materials and Methods

### Sample collection

Water and microbialite samples were collected in Great Salt Lake (Antelope Island State Park, Utah 41° N, 112° W, GSL, near Layton, Utah) and Green Lake (Green Lake State Park, New York 43.049°N, 75.973°W, FGL, near Fayetteville, New York).

### Preparation of nucleic acid stain working stock

The working stock was created from a commercial stock of SYBR Gold (Invitrogen S11494), thawed in the dark at room temperature. Once thawed, the commercial stock was vortexed for 10 s at medium-high speed, then centrifuged in a microcentrifuge for five minutes. The commercial stock was then diluted 1:10 with autoclaved and filtered (0.22 μm PVDF Millipore GVWP06225) molecular biology grade water. The working stock was filtered (0.22 μm PVDF filters) before small volumes (∼300 μL) were aliquoted into microcentrifuge tubes wrapped in aluminum foil or black microcentrifuge tubes and stored at -20°C until use.

### Preparation of samples

Samples were prepared similarly across the water, whole microbial mat, and EPS, which included filtration, differential centrifugation, chloroform treatment, glutaraldehyde fixation, benzonase nuclease treatment, probe sonication (EPS and mat only), SYBR Gold staining, wet-mounting, and imaging (**Fig 1**). Water samples from GSL and FGL were prepared with an optional concentration step using 30 kDa MWCO pore-size centrifugal filter devices (Millipore UFC5030). Water samples (i.e., 500 mL) were filtered twice through 0.22 μm PVDF filters and then concentrated in Centricon-70 plus centrifuge filters (Millipore UFC703008, 30 kDa) in 12 min increments at 3,500 RPM. Flow through water (i.e., ultrafiltrate) was collected after each centrifugation step. A final centrifugation step of 15 min at 3,500 RPM was used to collect the filtrate by flipping the filter into the collection cup to obtain a concentrated viral sample. Viral concentrated water was recovered from Centricon-70, then diluted to 1 mL with ultrafiltrate from the original location.

**Figure 1.**
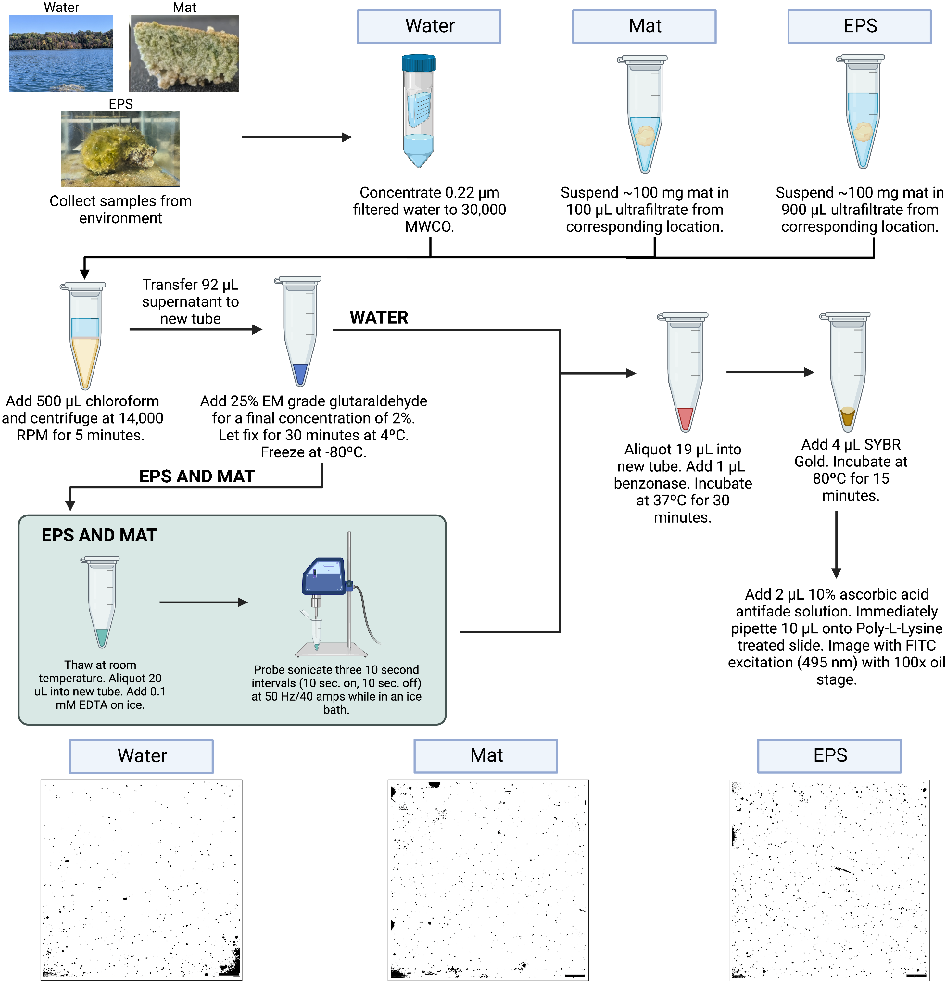
Flow diagram of EFM protocol. This diagram provides a detailed methodological walk-through of the preparation of water, microbial mat, and EPS samples.

Whole microbial mats and EPS were resuspended in matching location ultrafiltered water (i.e., 30 kDa filtered). Approximately 100 mg of the microbial mat was taken for the whole mat vs. EPS samples. The whole mat was resuspended in 100 μL of ultrafiltrate vs. 900 μL of ultrafiltrate for EPS (**Fig 1**). All samples were treated with 500 μL of chloroform, then differential centrifugation, fixation with glutaraldehyde (2% final concentration), benzonase treatment, staining with SYBR Gold, then wet mounting (**Fig 1**). EPS and mat samples required additional treatments to get rid of cell debris, FED, and MDVs. Before benzonase treatment, EDTA was added to EPS and the whole mat for a final concentration of 0.1 mM, as suggested by Carriera et al. (2015). Then, samples were probe sonicated, and the supernatant was transferred to a new tube prior to nuclease treatment by benzonase (**Fig 1**). For all samples, benzonase treatment included 19 μL of sample with 1 μL benzonase (25 U) added, then incubated in a heat block at 37ºC for 30 minutes (**Fig 1**). Samples were frozen and then stored at -80°C until use.

### Preparation of wet-mounting, slide preparation, and microscopy

Slides were prepared before thawing samples by thoroughly cleaning them with 70% ethanol. After cleaning, the slides were allowed to dry before soaking in a 10% poly-L-lysine solution in a polystyrene dish for five minutes. After soaking, the slides were drained and dried in a drying oven at 60°C for one hour. The slides were then removed from the oven and stored in a microscope slide box until use.

SYBR Gold stock was aliquoted in the dark to avoid stain decay, then 4 μL of SYBR Gold working stock was added to each sample and pipetted to mix. The tubes were then incubated at 80°C in a heat block for 15 min in the dark. After incubation, samples were removed from the heat block and mixed with 2 μL of freshly prepared ascorbic acid antifade solution (10% ascorbic acid w/v in 1x PBS and filtered through 0.22 μm PVDF filters twice) (Cunningham et al., 2015), 10 μL of each sample was immediately pipetted onto poly-l-lysine treated slides, covered with a cover slide, and imaged on an Olympus IX83 microscope under FITC blue excitation light (495 nm) with a 100x oil stage (**Fig 1**).

### Enumeration

VLPs were counted in EFM images according to size and shape. Particles were included if they were <300 nm, which can be trained through images of beads of known sizes (data not included). Large particles, halo autofluorescence, and clumps of VLPs were excluded from counting. At least three slides with at least three images per slide and a minimum of 12 images total were averaged for each by location and sample type (e.g., GSL mat) (**Fig 2**). Automatic counts from programs such as ImageJ provided inaccurate results. Due to this, all images were manually counted and corrected by eye. Images with fewer than 100 VLPs and more than 500 VLPs were excluded. Exposure and contrast were adjusted to increase countability, and images were converted to negatives to increase the visibility of small particles. The field of view was calculated by finding the image’s area in μm using the length and width found with the Olympus cellSens software. Cover slides of 22×22 mm were used. The area of the cover slides was calculated by multiplying the length by the width and was converted to μm. The scaling factor was calculated by dividing the cover glass area by the field of view. This value was then used to multiply the counts for each image to determine the number of viral particles in 10 μL used per slide. Water samples were adjusted to account for concentration, then converted to determine VLPs mL^-1^. VLP concentrations for microbial mat and EPS samples were calculated by multiplying the counts by the same scaling factor as with water samples, dividing by 10 μL, multiplying by the dilution factor, and then dividing by the mass of microbial mat or EPS sample used (∼0.10 g).

**Figure 2.**
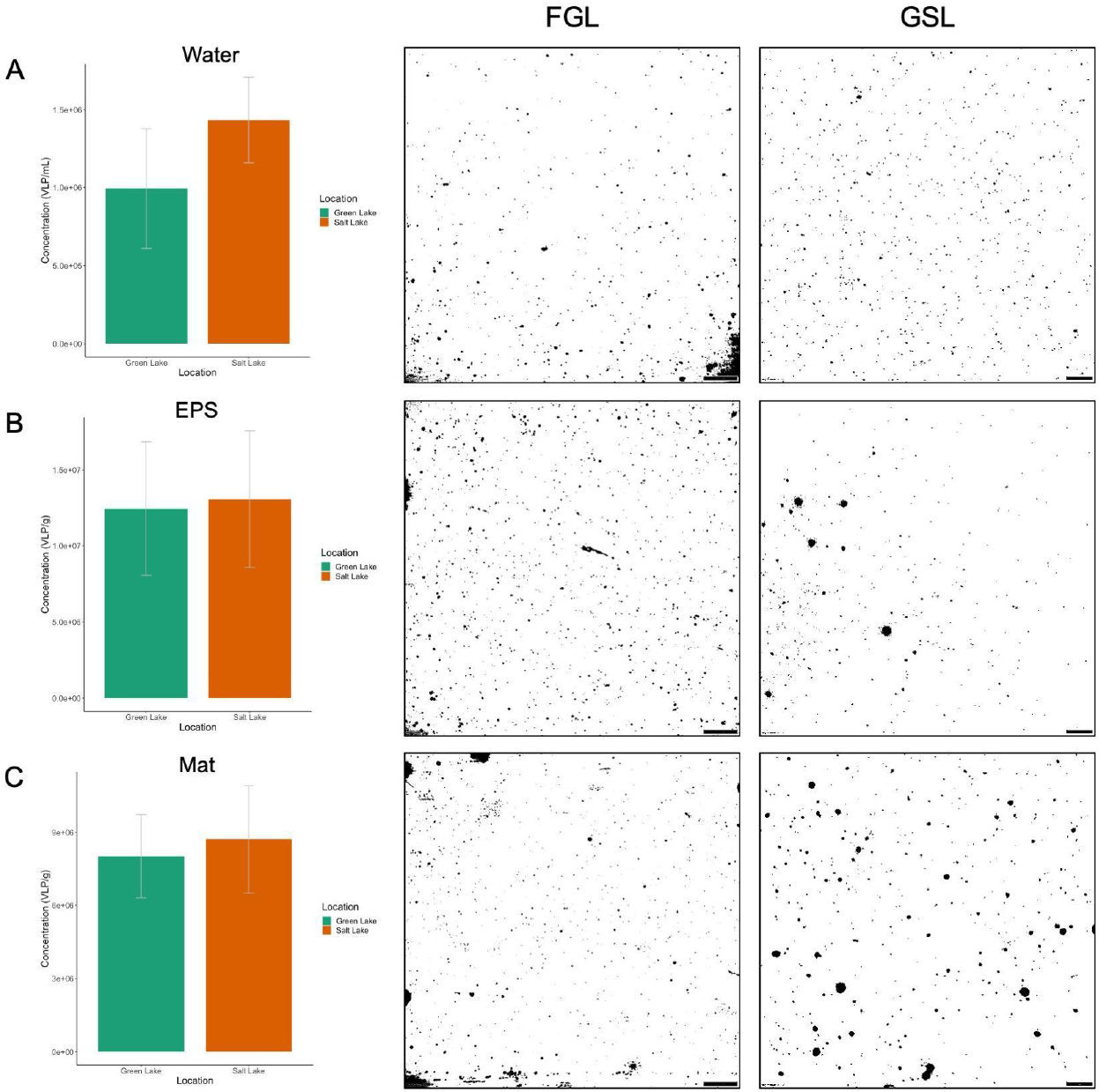
Viral Enumeration of GSL and FGL. EPS and whole mat VLPs are expressed as g^-1^ vs. mL^-1^ for water samples. All images are under a 100x oil immersion with a scale bar equal to 200 pixels or 10 µm. (A) VLP concentrations for water samples with representative EFM images. (B) VLP concentrations for EPS samples with representative EFM images. (C) VLP concentrations for mat samples with representative EFM images.

### Data Availability Statement

Raw image files and supplemental data are all available at https://osf.io/f8be4/. All code and calculations for this study are available on www.github.com/raw-lab/efm.

## 3. Results

VLPs across sample types varied but not across ecosystems (FGL vs. GSL). GSL and FGL had similar viral abundances at ∼1 × 10^6^ VLPs mL^-1^ (i.e., 1.43 × 10^6^ mL^-1^ GSL vs. 9.95 × 10^5^ VLPs mL^-1^ FGL, **Fig 2**). GSL VLPs appeared to have more uniform sizes and were smaller on average than FGL (**Fig 2**). The coefficient of variance (CV) was higher in FGL than in GSL (38.79% FGL vs. 19.15% GSL). Large particles were present in FGL water that were >300 nm and lacked uniform VLP size. EPS had the highest VLP counts observed. EPS had an order of magnitude higher abundance of VLPs than water ∼1 × 10^7^ VLPs g^-1^ (i.e., 1.25 × 10^7^ g^-1^ FGL vs. 1.31 × 10^7^ VLPs g^-1^ GSL) with similar CVs (35.26% FGL vs. 34.40% GSL, **Fig 2**). Whole mat samples had higher VLPs than water as well 8-9 × 10^6^ g^-1^ (i.e., 8.02 × 10^6^ g^-1^ FGL vs. 8.72 × 10^6^ VLPs g^-1^ GSL) with similar CVs (21.32% FGL vs. 25.33% GSL, **Fig 2**). GSL and FGL did not differ statistically across ecosystems within the sample type (e.g., EPS FGL vs. GSL) and had similar viral abundances. Low (< 4.0 × 10^5^ VLPs/mL) and high (> 2.0 × 10^7^ VLPs/mL) samples are well within our detection limits using this method.

Our protocol added a chloroform step to all samples and a benzonase treatment to aquatic samples. The chloroform and benzonase treatment did remove large cellular debris, MDVs, and FED (data not shown). We confirm the results from Carriera et al. (2015) that probe sonication is superior to water bath-based, EDTA was better than tetrasodium pyrophosphate, and that benzonase treatment cleared FED better than DNase I only or DNase/RNase (data not shown). Chloroform offered greater removal of cellular debris of aggregated VLPs or MDVs in carbonate-rich EPS and mat from microbialites than without (data not shown).

The Cunningham method requires chemical flocculation concentration of the VLPs using iron chloride precipitation, followed by EDTA-ascorbate resuspension for preparing samples (Cunningham et al., 2015). We tested this on our aquatic samples and had similar results but found that chloroform and benzonase treatment cleared the images providing better quality. We found high concordance in aquatic samples with and without chemical flocculation. Thus, we opted out of using it due to the extra steps required. Chemical flocculation did not work for EPS or mat from modern microbialites in our hands. We were unable to resuspend samples post-flocculation.

Poly-l-lysine was another addition to our protocol over others that provided an added benefit. The addition decreased CVs across images, allowed for more precise placement of VLPs across the slide, made the field of view of VLPs more uniform, and provided greater storage stability of VLPs on slides. Slides made even a month later had <1% drop in viral counts using poly-l-lysine (data not shown).

Cost analysis suggests that Anodisc filter-based methods are >$15 dollars a sample (**Table 1**). The Cunningham method is currently the most cost-effective method, originally costing ∼$0.10 per sample in 2015, but the original beads used in his study are no longer in manufacture (**Table 1**). We used a similar-sized bead that is available for our cost estimation (2.06 µm bead) which, due to inflation, now makes the method cost almost double in less than a decade (∼$0.18 per sample) (**Table 1**). We recommend benzonase and chloroform treatment with the standard Cunnigham method; the price increases to ∼$0.84 per sample, but both will improve image quality (**Table 1**). Our method is ∼$0.75 per sample, mainly due to the cost of benzonase, which is more expensive than DNase I/RNase I combined but is superior (**Table 1**).

**Table 1.** Cost analysis of EFM approaches* for counting bacteria and viruses in water, microbial mats, and EPS. The cost of consumables (pipette tips, microcentrifuge tubes, etc.) is estimated to be the same for each method.

| Method | Cost (USD) |
| --- | --- |
| Bellanger et al., presented here | \$0.75 |
| Cunningham et al., 2015 (water) | \$0.18** |
| Advanced Cunningham (water) | \$0.84*** |
| Carreira et al., 2015 (mat) | \$18.79 |
| Budinoff et al., 2011 (water) | \$22.12 |
| Noble and Fuhrman, 1998 (water) | \$16.18 |

\*Values calculated during the COVID-19 pandemic and inflation period of 2022 have caused costs and supply fluctuations.
\*\*Original bead size (2.34 $\mu\text{m}$ ) is no longer in manufacture. A similar size bead (2.06 $\mu\text{m}$ ) was used to calculate cost instead.
\*\*\*Advanced method is the Cunningham method with adding chloroform and benzonase.

## 4. Discussion

Our EFM protocol is an integration of multiple methods in order to provide results across both aquatic and solid substrate ecosystems (Forterre et al., 2013; Carreira et al., 2015; Cunningham et al., 2015). These integrations are used to analyze VLPs over fake particles (Forterre et al., 2013). Our study confirms suggestions that chloroform and nuclease treatment clears images and removes FEDs, MVDs, and cellular debris. Furthermore, it confirms the results of Carriera et al. (2015) with some additions that can be applied to microbial mats and EPS on solid carbonate microbialites, including a wet-mount method that removes the expensive Anodisc filter requirement. We could not use chemical flocculation on microbial mats and EPS from modern microbialites with any success. Further method development is needed to use chemical flocculation within microbial mats and EPS in modern microbialites or other solid substrate ecosystems such as soils or sediments.

FGL and GSL had relatively the same VLP concentrations regardless of the sample type and did not differ statistically. EPS had the highest VLP concentration at ∼10^7^ g^-1^. However, the whole microbial mat also had similar abundances 8-9 × 10^6^ g^-1^, which was an order of magnitude higher than surrounding water suggesting VLPs are being trapped or concentrated within the mat and/or EPS. Observation of whole microbial mats shows VLPs trapped on the surface layers, which has been suggested previously (Carriera et al., 2015; Pacton et al., 2014), which we confirm here. These viruses may act as nucleation sites for carbonate minerals within the microbialite mat (Pacton et al., 2014; Słowakiewicz et al., 2021).

Previously, soft intertidal marine microbial mats (i.e., not hard carbonate microbialites) in Schiermonnikoog Island, the Netherlands, had some of the highest measured VLPs ever recorded at 2.8 × 10^10^ g^-1^ (Carriera et al., 2015). We find that GSL and FGL microbialite mats do not have these similar high abundances as intertidal marine microbial mats. They are more similar to standard aquatic VLP abundances within freshwater samples ∼10^6^ to 10^7^. EPS within GSL and FGL microbialites are in the low range of VLPs abundance in soils which ranges from 10^7^ to 10^9^ (Williamson et al., 2017). Our microbialites are present in meromictic lakes that do not seasonally turnover or mix beyond rainfall, suggesting less bacterial mixing, whereas intertidal zones have continuous mixing and influx of microbes, including virally infected microbes. Previous metagenomic analysis of both Pavilion Lake and Shark Bay microbialites and microbial mats found viruses present, but high gene abundance of viral defense clusters (White III et al., 2016; White III et al., 2018; Wong et al., 2018). Pavilion Lake metagenomes had statistically higher viral genes present in microbialite surrounding water metagenomes than microbialite metagenomes (White III et al., 2016).

Viral counting is done manually by eye for all current EFM methods. While software such as ImageJ exists; however, it struggles with automatic counts within images with large particles, halo autofluorescence, and clumps of VLPs. ImageJ was unable to count VLPs accurately across our samples in general. While counting manually is very labor intensive and requires training for accurate VLP counting, it is our only current option. Manual counting may miss dim low fluorescence containing VLPs, small VLPs, and VLPs underneath halo autofluorescence, and general human error can occur, but it is limited by proper training. Future software and algorithm development is needed to alleviate the labor-intensive nature of manual VLP counting for EFM.

Currently, our method enumerates viruses regardless of the nucleic acid type, strandedness and accounts for the large viruses. Further improvements are needed in EFM dyes to distinguish single-stranded vs. double-stranded nucleic acids when mixed. Also, giant viruses, including pandoraviruses, are commonly filtered out during standard filtration of 0.22 μm (Philippe et al., 2013). Acridine orange may be selective for RNA over DNA in fluorescence shift and effective for ssDNA phages (Darzynkiewicz, 1990; Mayor & Hill, 1961). Chloroform without filtration has been useful in removing bacteria to isolate jumbo and megaphages (Saad et al., 2019). Antibiotics and fungicides may be useful for membrane-bound giant viruses like pandoraviruses to remove bacteria and fungi in enumeration studies (Philippe et al., 2013). DAPI and Yo-Pro-I were used for viral enumeration but were too dim compared to SYBR-based options (Patel et al., 2007). SYBR Green I and SYBR Gold were equivalent for virus and bacterial enumeration (Patel et al., 2007). SYBR green I/II or SYBR gold can be used for RNA virus staining (Patel et al., 2007), but not in RNA and DNA virus mixed samples.

Counts of viruses in EFM samples can broadly estimate the abundance of viruses within the water, microbial mats, and overall systems of FGL and GSL. The lake volumes can change for various environmental (i.e., seasonally, precipitation) and anthropogenic reasons. Similarly, viral concentrations may vary seasonally and spatially (with depth and relation to the shore) (Brum et al., 2016). With these understandings, approximations of viral abundance and total weight are derived from our measurements. The volume of FGL is estimated to be 7,235,900 m^3^ (Brunskill & Ludlam, 1969). For FGL, based on the total volume, which is 7.2 × 10^12^ mL times the average VLP concentration of 9.95 × 10^5^ mL^-1^, the total number of viruses is 7.2 × 10^18^ VLPs. If we assume that each virus weighs at least 50 × 10^6^ daltons (based on coliphage T7) (Rontó et al., 1983) and convert daltons to grams using the Avogadro constant (1.66 × 10^−24^), then the weight of all the viruses present within FGL is ∼598 g. A standard loaf of bread weighs roughly 500 g. The volume of all FGL viruses, if concentrated all together, could fit in a standard salad dressing bottle. GSL is estimated to contain 11.42 to 18.92 km^3^ (Baskin, 2005). Converting km^3^ into mL yields 1.1-1.9 × 10^16^ mL total volume times the average VLP concentration of 1.43 × 10^6^ mL^-1^, results in ∼2.7 × 10^22^ VLPs total in GSL. If we assume that the average weight is 50 × 10^6^ daltons, and convert daltons into kilograms using the Avogadro constant (1.66 × 10^−27^), then the weight of all the viruses in GSL is ∼2.2 kg. Four footballs, twelve apples, and a standard-size brick are each ∼2 kg. A standard gallon milk jug would fit all the viruses from the Great Salt Lake within a standard refrigerator.

Further advancements are needed to directly measure viruses to understand how many viruses are present within natural ecosystems beyond what is presented here. The development of EFM methods using various nucleic acid, protein, and lipid stains may help to distinguish between intact viral particles and expose their presence regardless of their nucleic acid type, strandedness, or size. Seasonal measures of the viral abundances are needed within GSL and FGL waters and microbialites to enumerate any variation, especially GSL, which has experienced record lows due to droughts. Enumeration is the first critical step in the ecology of an ecosystem. Our method is an advancement towards robust viral measurement within modern microbialites, microbial mats, and aquatic ecosystems that is cost-effective (∼$0.75 per sample).

## Acknowledgments

We also acknowledge the University Research Computing and the College of Computing and Informatics for computational and logistical support.

## Author Contributions

Conceptualization, R.A.W. III, M.A.B., and P.V.; Methodology, M.B. and R.A.W. III; Investigation, M.A.B. and R.A.W. III,; Writing – Original Draft, M.A.B. and R.A.W. III,; Writing - Review & Editing, R.A.W. III, M.A.B., and P.V.; Funding Acquisition, R.A.W. III and P.V.; Resources, R.A.W. III and P.V.; Supervision, R.A.W. III.

